# ISWI chromatin remodeler SMARCA5 is essential for meiotic gene expression and male fertility in mammals

**DOI:** 10.1101/2024.10.14.618292

**Authors:** Shubhangini Kataruka, Aushaq B Malla, Shannon R Rainsford, Bluma J Lesch

**Affiliations:** Department of Genetics, Yale School of Medicine, New Haven CT USA 06510; Department of Obstetrics, Gynecology & Reproductive Sciences, Yale School of Medicine, New Haven CT USA 06510; Yale Cancer Center, Yale School of Medicine, New Haven CT USA 06510

## Abstract

Regulation of the transcriptome to promote meiosis is important for sperm development and fertility. However, how chromatin remodeling directs the transcriptome during meiosis in male germ cells is largely unknown. Here, we demonstrate that the ISWI family ATP-dependent chromatin remodeling factor SMARCA5 (SNF2H) plays a critical role in regulating meiotic prophase progression during spermatogenesis. Males with germ cell-specific depletion of SMARCA5 are infertile and unable to form sperm. Loss of *Smarca5* results in failure of meiotic progression with abnormal spermatocytes beginning at the pachytene stage and an aberrant global increase in chromatin accessibility, especially at genes important for meiotic prophase.

## Introduction

Meiotic cell division is a unique process exclusive to germ cells and essential for production of haploid gametes. Meiotic entry, progression and exit are actively regulated processes, and their failure leads to subfertility or sterility. Regulation of meiotic progression requires a specialized epigenome and is supported by extensive chromatin remodelling before, during and after meiosis in both male and female germ cells. Epigenome remodeling during meiosis is important for transcription of meiotic regulatory genes, suppression of retrotransposons that can threaten genomic integrity, and homologous recombination required for proper chromosome segregation and generation of genetic diversity (Peters et al. 2001; Tachibana et al. 2007; Sasaki and Matsui 2008). For example, the genome-wide deposition pattern of the histone modification H3K9me2 changes dynamically during the transition from pre-meiotic spermatogonia to primary spermatocytes in prophase I of meiosis, and knockout of the H3K9 methyltransferases *Suv39h1* or *Suv39h2* correspondingly leads to abnormal meiotic prophase in males. DNA methylation and small RNA pathways are also actively regulated, well characterized epigenetic mechanisms necessary for silencing of retrotransposons during meiosis, and perturbation of either DNA methylation or the piRNA pathway leads to male sterility (Bourc’his and Bestor 2004; Aravin et al. 2007). However, the complete set of mechanisms that regulate the complex chromatin remodelling required for progression through meiosis is not yet defined.

ATP-dependent chromatin remodelers can activate expression of stage-specific genes while supressing inappropriate transcription during development by altering interactions between histones and DNA. There are four classes of ATP dependent remodelers: SWI/SNF (switch/sucrose non-fermentable), ISWI (imitation switch), CHD (chromodomain helicase DNA-binding) and INO80 (SWI2/SNF2 related (SWR)), all of which share a similar ATPase domain (Cote et al. 1994). SWI/SNF, CHD, and INO80 family members have all been shown to play important roles in spermatogenesis and early embryogenesis, but the role of the ISWI remodelers in spermatogenesis is unknown (Bultman et al. 2006), Kim, Fedoriw et al. 2012, Li, Wu et al. 2014), (O’Shaughnessy-Kirwan et al. 2015; Suzuki et al. 2015), (Wang et al. 2014; Serber et al. 2016).

Mammals have two ISWI paralogs, *Smarca1* (*Snf2l*) and *Smarca5* (*Snf2h*), which have distinct expression profiles and functions (Lazzaro and Picketts 2001). Deletion of *Smarca1* has no discernible phenotype in adult mice (Yip et al. 2012), whereas *Smarca5* is essential for survival beginning in the earliest stages of embryogenesis. Oocyte-specific conditional deletion of *Smarca5* prevents ovulation due to failure to re-enter meiosis following arrest during fetal stages, and female germline conditional knockout mice are sterile (Zhang et al. 2020). Recently, *Smarca5* has also been shown to be an important regulator of zygotic genome activation (ZGA) in mice (Oana Nicoleta Kubinyecz 2023), and *Smarca5* knockout embryos show peri-implantation lethality because of failure to proliferate in both the inner cell mass and trophectoderm (Stopka and Skoultchi 2003). We hypothesized that SMARCA5 could play an important role in male meiosis similar to its requirement for meiotic progression in females.

Here, we conditionally deleted *Smarca5* specifically in mouse male germ cells at two stages of spermatogenesis, and found that *Smarca5* is essential for male meiotic progression. Loss of *Smarca5* from male germ cells leads to sterility, accumulation of aberrant pachytene-like cells, reduced numbers of post-meiotic round spermatids, and complete absence of elongated spermatids and epididymal sperm. Cells that have entered meiosis in *Smarca5* cKO testes have increased rates of apoptosis and elevated expression of LINE1 retrotransposons. Single-cell transcriptomics confirmed the accumulation of an abnormal pachytene-like cell population, concommitant with extensive transcriptional misregulation. At the chromatin level, *Smarca5* cKO germ cells exhibited extensive gains in chromatin accessibility, including at genes whose proper regulation is essential for meiotic progression. This effect is consistent with previous reports that ISWI remodellers act to compact chromatin but contrasts with the reported role of SMARCA5 in promoting chromatin opening and gene activation during oocyte meiosis. Therefore, even though *Smarca5* is essential for promoting meiotic gene expression and meiotic progression in both male and female germ cells, its effects differ between the sexes, reflecting different regulatory requirements in male and female meiosis. Together, our results reveal that SMARCA5 facilitates germ cell progression through male meiosis by coordinating proper chromatin accessibility and transcriptional regulation required for successful gamete development.

## Results and Discussion

To address the role of SMARCA5 in spermatogenesis, we generated a germ cell-specific knockout of *Smarca5* (*Smarca5* cKO) by crossing mice carrying a conditional allele of *Smarca5* (*Smarca5*^*fl/fl*^) (Alvarez-Saavedra et al. 2014) with mice carrying the *Ddx4-Cre* allele (Gallardo et al. 2007; Hu et al. 2013). Expression of *Ddx4-Cre* begins around E10.5-11.5 in both male and female germ cells and continues in the germ cell lineage throughout life, with excision of the conditional allele complete by the time of birth. *Smarca5* cKO mice are thus expected to express only a truncated SMARCA5 protein without the ATPase domain at all stages of spermatogenesis and all postnatal ages (**Supplemental Fig S1A-B**). Western blot and immunostaining in testis confirmed a significant reduction of SMARCA5 protein in adult mouse testis, with residual signal in the Western blot likely explained by the somatic cells present in whole-testis samples (**Fig 1A, 1B**). Immunostaining showed strongest expression in spermatogonia and early-stage spermatocytes at the periphery of the tubules, consistent with previous reports that SMARCA5 protein expression is highest in spermatocytes (**Fig 1B**) (Chong et al. 2007). We further confirmed that the phenotype in *Ddx4-Cre* cKO females matched that of the previously reported *Zp3-Cre*-driven *Smarca5* conditional knockout females, which were infertile due to lack of ovulated oocytes (**Supplemental Fig S1C**) (Zhang et al. 2020). We conclude that our *Smarca5* cKO mice appropriately model loss of SMARCA5 in male germ cells.

**Figure 1.**
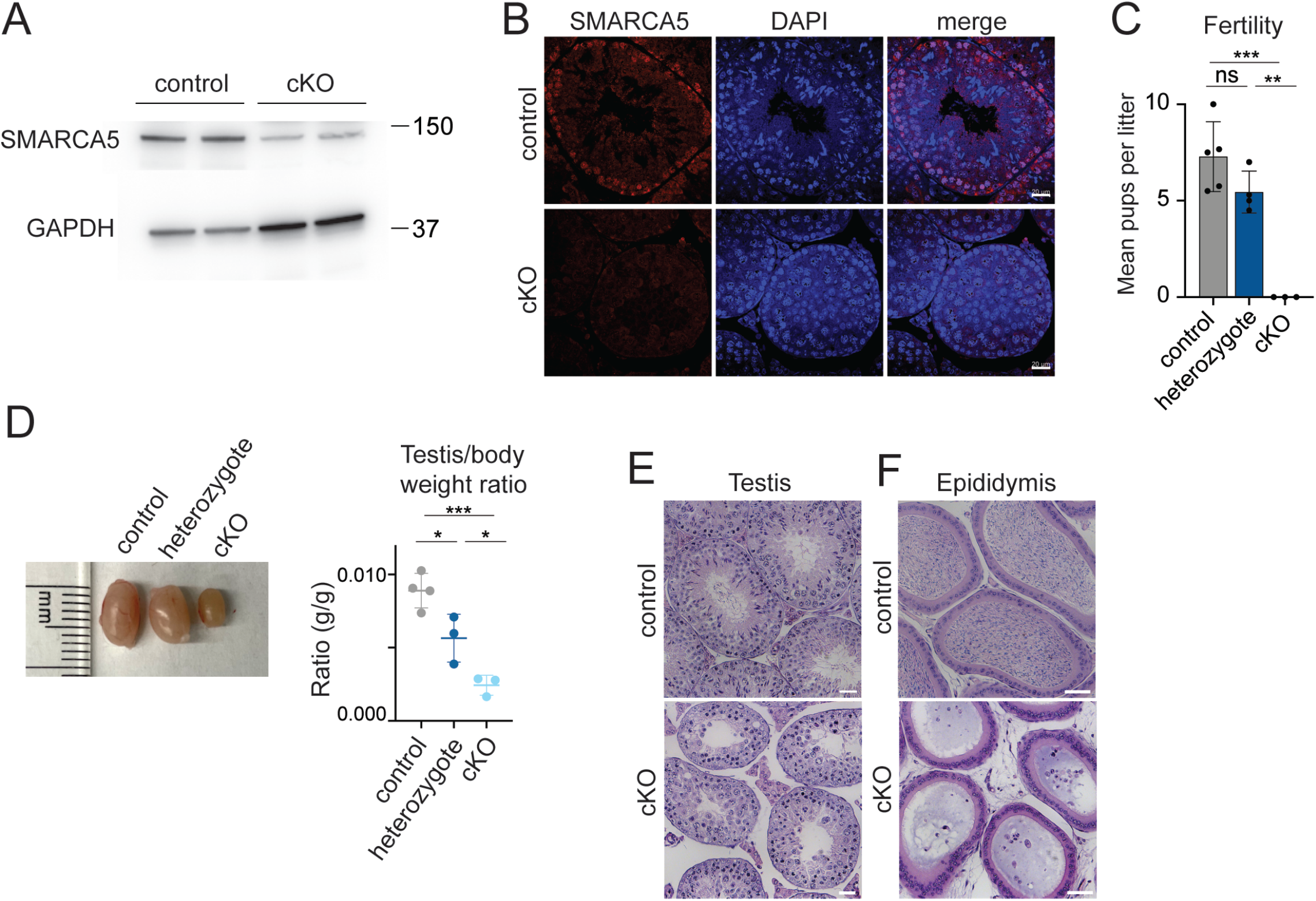
*Smarca5* is essential for male fertility in mouse. **A**, Western blot for SMARCA5 from wild type and *Smarca5* conditional knockout (cKO) adult whole testes. GAPDH is a loading control. **B**, Immunostaining of SMARCA5 in paraffin sections from control and *Smarca5* cKO adult testes. DNA is stained with DAPI. Scale bar, 20µm. **C**, Fertility test for wild type, *Smarca5* heterozygote, and *Smarca5* cKO (n=3-5). Each dot represents mean litter size for one male. Error bars represent standard deviation. No pups were born from *Smarca5* cKO males. *p<0.05, **p<0.01, ***p<0.01, unpaired Welch’s t-test. **D**, Left, gross image of whole testis isolated from control, heterozygous, and *Smarca5* cKO adult male mice. Each tick on the ruler represents 1mm. Right, testicular weight relative to body weight plotted for n=3-4 control, heterozygous, and *Smarca5* cKO testes. *p<0.05 **p<0.01 ***p<0.001, unpaired t-test. **E-F**, Hematoxylin and eosin (H&E) staining of paraffin-embedded sections from control and *Smarca5* cKO adult testis (**E**) and epididymis (**F**). Scale bars, 100µm.

We found that *Smarca5* cKO male mice are infertile, as no pups were born when they were allowed to mate freely with wild type female mice for six months (**Fig 1C**). Testes of *Smarca5* cKO mice were much smaller than controls (**Fig 1D**). By histology, there were very few round spermatids and almost no elongating spermatids and mature sperm in the seminiferous tubules, and epididymides were devoid of sperm (**Fig 1E, 1F**). Interestingly, we also observed that heterozygous *Smarca5* male mice were subfertile and had testis-to-body-weight ratios intermediate between wild type and cKO, suggesting dosage sensitivity (**Fig 1C, 1D**). We conclude that SMARCA5 is required for male fertility and normal spermatogenesis.

To further support the defects observed by histology, we quantified the proportions of cells at different spermatogenic stages by flow cytometry, using the nucleic acid dye propidium iodide (PI) to detect populations of cells with different chromosomal complements. In agreement with the absence of elongated spermatids and mature sperm seen by histology, there was a significant reduction in the fraction of elongating spermatids and a trend toward reduced round spermatids in *Smarca5* cKO testes, suggesting a defect during or before meiosis (**Fig. 2A**). Conversely, there was a significant increase in the spermatogonia and spermatocyte populations, suggesting an abnormal accumulation of cells due to failure to progress through meiotic prophase. *Smarca5* cKO testes showed evidence of multiple defects that could contribute to failure of meiotic progression, including elevated levels of the DNA damage marker γH2A.X (**Fig 2B, Supplemental Fig S1D**), higher numbers of apoptotic cells as assayed by TUNEL (**Fig 2C**), and higher expression of LINE1 protein indicating aberrant activation of transposable elements (**Fig. 2D**). De-repression of transposable elements could be the direct cause of germ cell loss and infertility, or could be an indirect effect of changes in epigenetic state caused by the loss of SMARCA5 chromatin remodeling activity.

**Figure 2.**
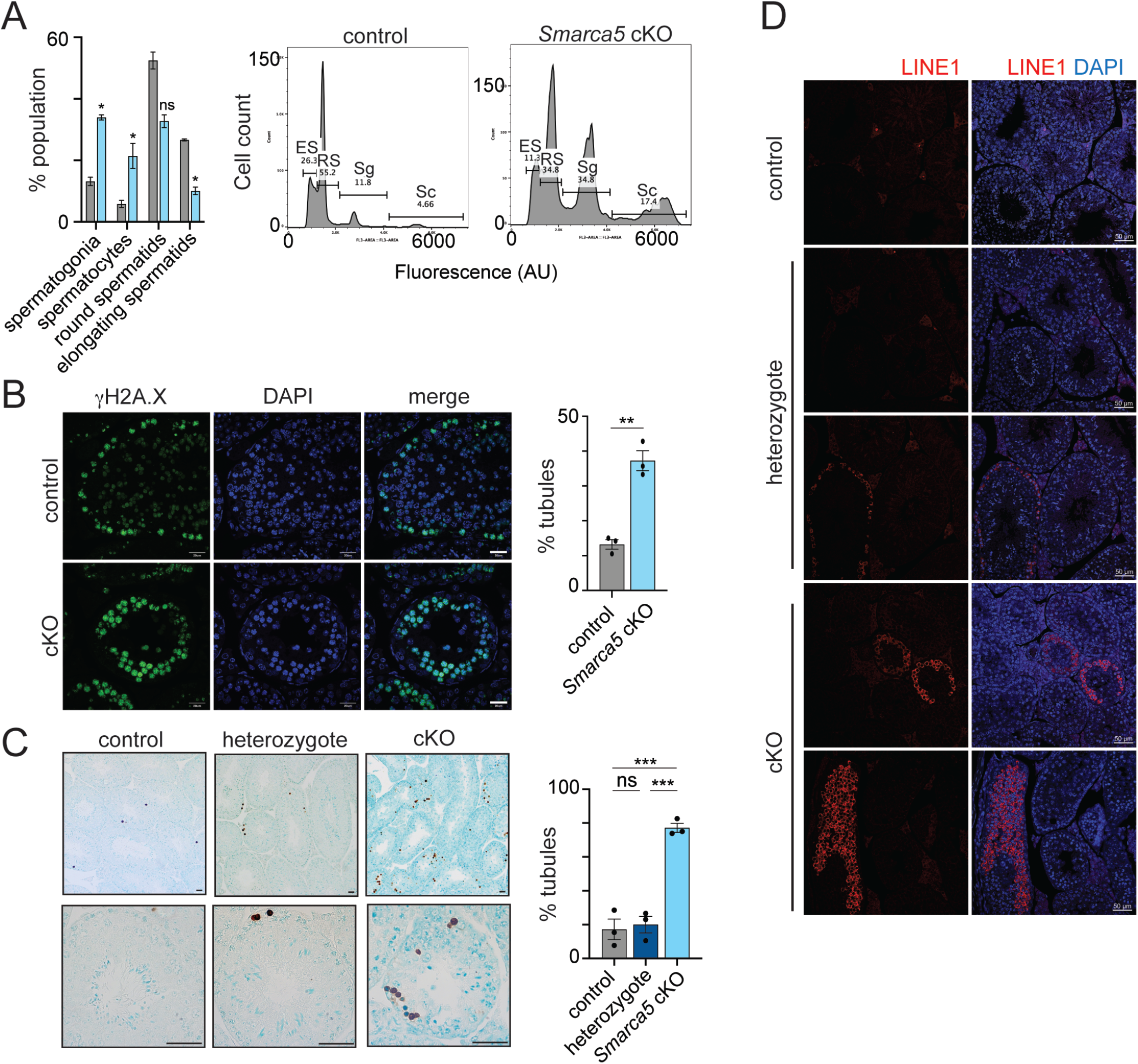
Deletion of *Smarca5* results in defective meiotic progression and LINE1 derepression. **A**, Flow cytometry for testicular cells from wild type or *Smarca5* cKO males stained with propidium iodide to measure DNA content. Sg, spermatogonia; Sc, spermatocytes; RS, round spermatids; ES, elongating spermatids. **B**, Left, immunostaining of γH2AX in paraffin sections from wild type and *Smarca5* cKO adult testes. DNA is stained with DAPI. Scale bar, 20µm. Right, Fraction of tubules with nucleus-wide γH2AX staining across n=3 animals. **C**, Left, TUNEL assay for wild type, *Smarca5* heterozygous and *Smarca5* cKO adult testis paraffin sections. Right, fraction of TUNEL+ tubules across n=3 animals. Scale bar, 200µm. **D**, Immunostaining for LINE1 ORF1 protein in adult testis from control, heterozygote, and *Smarca5* cKO. DNA is stained with DAPI. Scale bar, 20µm. For all panels, *p<0.05, **p<0.01, ***p<0.01, unpaired t-test.

To better understand the timing and nature of meiotic defects induced by loss of SMARCA5, we prepared meiotic spreads from control, heterozygous, and *Smarca5* cKO testes. We confirmed that SMARCA5 expression is normally present in nuclei of control and heterozygous spermatocytes, and that its expression was severely depleted or absent in *Smarca5* cKO cells (**Fig 3A**). To evaluate when defects arise during meiosis in the absence of SMARCA5, we examined spreads co-stained for the synaptonemal complex protein SYCP1 and the DNA damage marker γH2A.X. At the leptotene stage, γH2A.X is ordinarily widespread due to the presence of naturally induced double-strand DNA breaks, and SYCP1 is beginning to assemble in linear patches at the chromosomal axes. We found that control, heterozygous and *Smarca5* cKO testes all contain similar numbers of normal-appearing leptotene spermatocytes based on SYCP1 and γH2A.X localization, indicating that *Smarca5* cKO cells can initiate meiotic prophase (**Fig 3B**). However, spreads from *Smarca5* cKO testes had few normal-appearing pachytene nuclei. Ordinarily, at the pachytene stage SYCP1 is fully assembled along the chromosomal axes and γH2A.X is restricted to the X or Y chromosome (the sex body). About half of *Smarca5* cKO pachytene-like cells have normal-appearing SYCP1 assembly along the chromosomal axes, but ectopic retention of gH2A.X outside of the sex body, indicating failure to fully resolve double-strand breaks during homologous recombination and repair (**Fig. 3C**, third row). In the other half of cKO pachytene-like cells, γH2A.X was appropriately restricted to the sex body but SYCP1 failed to assemble correctly on the chromosomes (**Fig 3C**, fourth row). Together, these data indicate that spermatocytes can initiate meiosis in the absence of SMARCA5, but fail to progress appropriately through the pachytene stage due to defective homolog pairing and repair of chromosome breaks.

**Figure 3.**
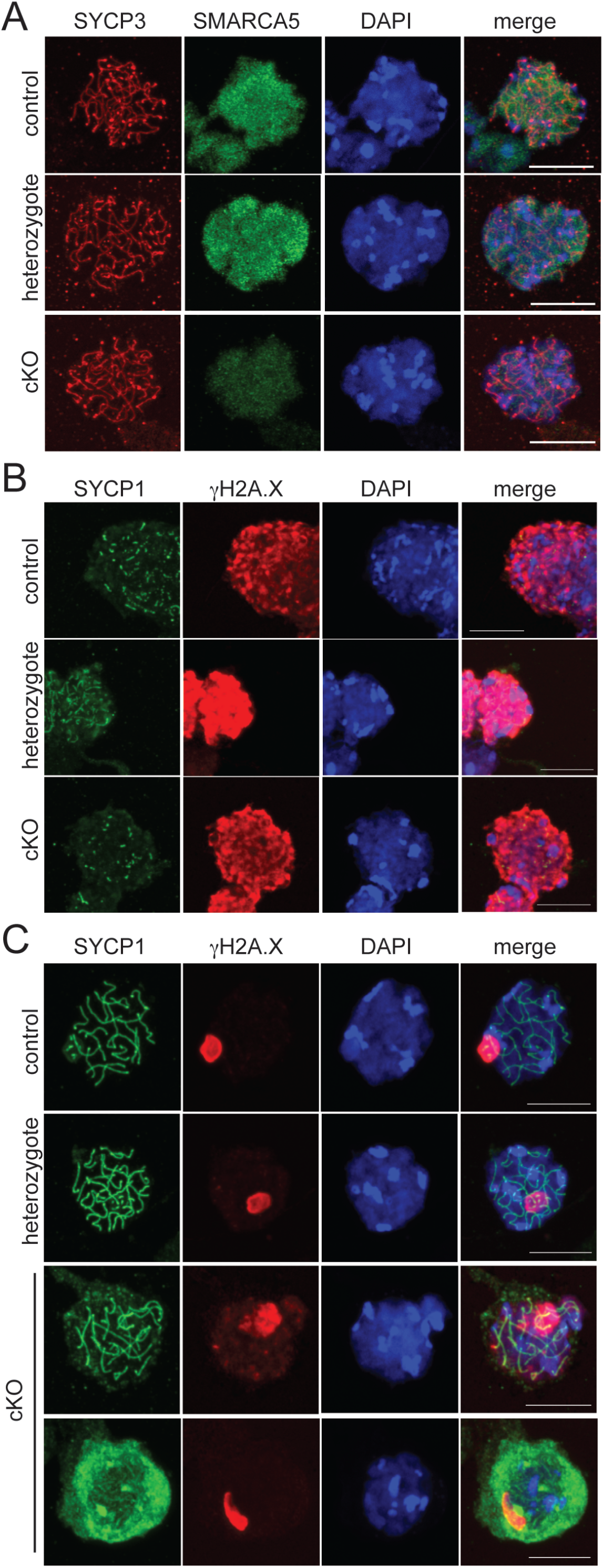
Spermatocytes lacking *Smarca5* exhibit defective chromosome synapsis in meiotic prophase I. **A**, Immunofluorescence staining of SMARCA5 and the synaptonemal complex marker SYCP3 in meiotic spreads from wild type, *Smarca5* heterozygous and *Smarca5* cKO adult testes. DNA is stained with DAPI. Scale bar, 10µm. **B**, Immunofluorescence staining of synaptonemal complex marker SYCP1 and DNA damage marker γH2AX in leptotene spermatocytes from wild type, *Smarca5* heterozygous and *Smarca5* cKO adult testes. DNA is stained with DAPI. Scale bar, 20µm. **C**, Immunofluorescence staining of SYCP1 and γH2AX in pachytene and pachytene-like spermatocytes from wild type, *Smarca5* heterozygous and *Smarca5* cKO adult testes. DNA is stained with DAPI. Scale bar, 20µm. For all panels, each image shows a single nucleus.

It is possible that meiotic defects observed using the early-acting *Ddx4-Cre* are secondary to earlier developmental defects in spermatogonial differentiation. To address this possibility, we generated a later deletion of *Smarca5* using the same *Smarca5*^*fl/fl*^ allele and a *Spo11-Cre* (*Smarca5;Spo11-Cre* cKO, **Supplemental Fig S2A**) (Lyndaker et al. 2013). *Spo11-Cre* is expressed in spermatocytes beginning at the very early stages of meiotic prophase, allowing for normal expression of SMARCA5 during spermatogonial differentiation. *Smarca5*;*Spo11-Cre* cKO mice were also sterile, with much smaller testes and a lower testis-to-body-weight ratio compared to control (**Supplemental Fig S2B, S2C**). Like the *Ddx4-Cre* cKO, *Smarca5;Spo11-Cre* cKO male testis also lacked elongated spermatids and mature sperm by histology, and the epididymis was devoid of sperm (**Supplemental Fig S2D, S2E**). *Smarca5;Spo11-Cre* cKO testes also displayed elevated levels of gH2A.X (**Supplemental Fig S2F**). The absence of elongated spermatids and mature sperm in *Smarca5;Spo11-Cre* cKO males solidifies our original observation that SMARCA5 is essential for appropriate progression of meiotic prophase in the mammalian male germ line. Notably, the *Smarca5;Spo11-Cre* cKO phenotype was not completely identical to the *Ddx4-Cre* phenotype: *Smarca5;Spo11-Cre* cKO testes contained more round spermatids, and there was no phenotype evident in heterozygotes (**Supplemental Fig S2C, S2D**). These differences may be due to the persistence of translated protein and *Smarca5* RNA carried over from spermatogonia before *Smarca5* deletion occurs in the *Spo11-Cre* cKO, or may reflect a separate role for SMARCA5 in spermatogonial development.

SMARCA5 is an ATPase-dependent chromatin regulator, so its loss is expected to result in altered chromatin configuration and transcriptional changes at target genes. To understand the transcriptomic changes that occur due to loss of SMARCA5 in male germ cells, we performed 10x single cell RNA sequencing (scRNA-seq) in *Smarca5* cKO testes and compared these data to an existing control dataset collected under identical conditions (GSE216343). Following filtering and harmonization, the final dataset included 7866 control and 7850 cKO cells distributed into 18 cell clusters (**Fig 4A and Supplemental Fig S3A-D**). Cluster identities were assigned based on established gene expression markers for spermatogenic populations (Green et al. 2018; Hermann et al. 2018; Lukassen et al. 2018) (**Supplemental Fig S3E**). Comparison of cell numbers across clusters between the two conditions was largely consistent with the cell population differences observed by histology and flow cytometry: numbers of premeiotic spermatogonia were equivalent, while postmeiotic elongating spermatids were strongly depleted in the *Smarca5* cKO (**Fig 4B, 4C**). As expected, there was an accumulation of cells assigned to early and mid-pachytene spermatocyte clusters in *Smarca5* cKO testes, supporting the conclusion that cKO cells fail to progress or progress more slowly through this stage. Interestingly, two overlapping clusters were defined at mid-pachytene, where one (“mid-pachytene A”) was strongly enriched in the *Smarca5* cKO condition and the other (“mid-pachytene B”) was present only in control. We interpret this to mean that *Smarca5* cKO spermatocytes do not attain a normal pachytene-like expression state but instead stall in an alternative, abnormal pachytene state. Meanwhile, a subset of cells continue to progress transcriptionally but cannot develop into normal sperm.

**Figure 4.**
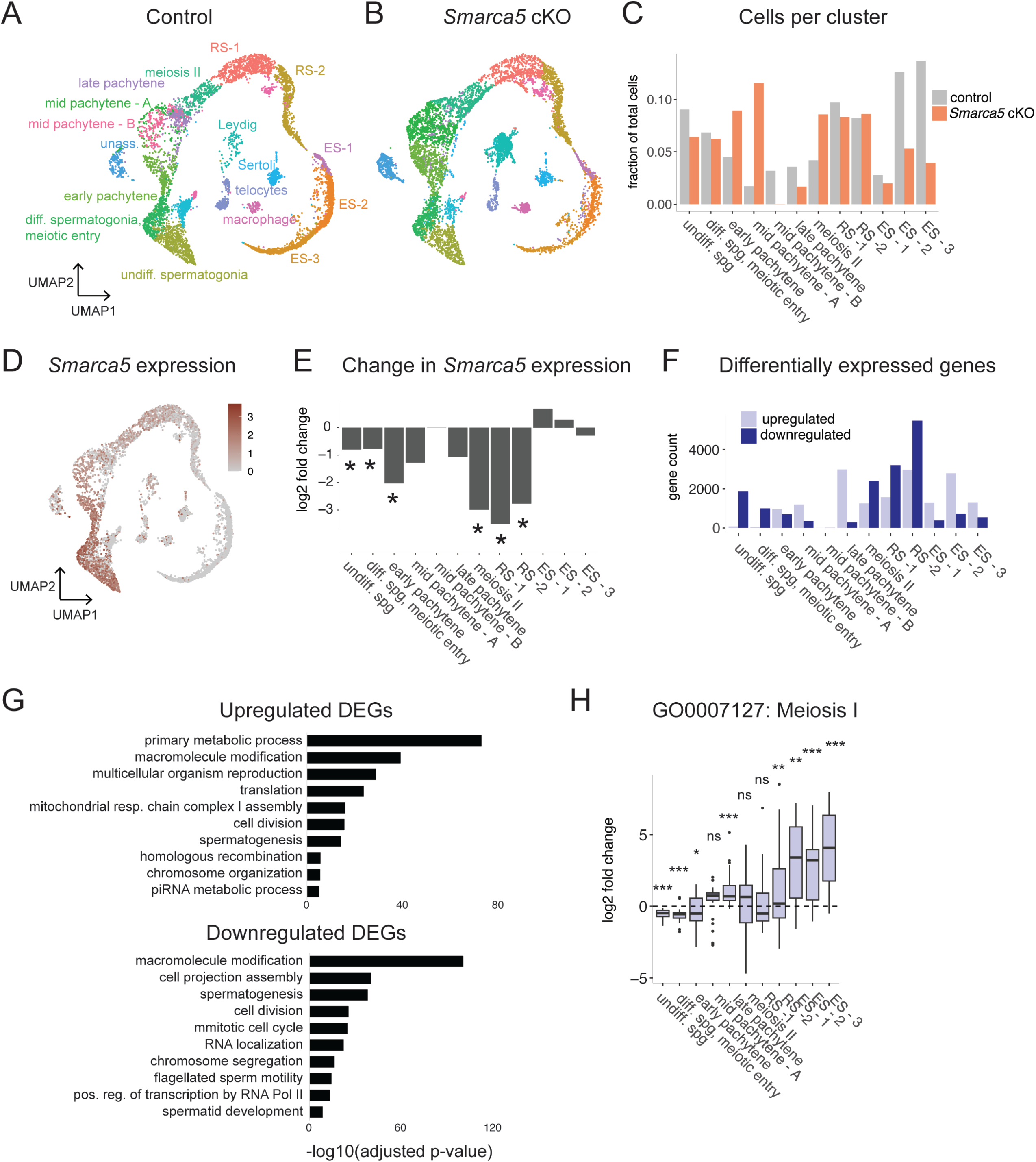
Transcriptomic dysregulation and accumulation of abnormal prophase I cells in the absence of SMARCA5. **A**, UMAP plot in control adult testes showing normal progression through meiosis and spermatogenesis. **B**, UMAP plot showing the same clusters in *Smarca5* cKO adult testes. **C**, Fraction of total cells in each cluster. The mid-pachytene – B cluster has zero cells in the cKO condition. Spg, spermatogonia. **D**, Expression of the *Smarca5* transcript projected on the wild type UMAP plot. **E**, Change in *Smarca5* expression in each cluster. *p<0.05 with correction for multiple hypothesis testing. **F**, Numbers of significant (p<0.05 after multiple testing correction) differentially expressed genes (DEGs) between wild type and *Smarca5* cKO in each cluster. **G**, Selected enriched Gene Ontology categories among upregulated and downregulated differentially expressed genes. **H**, Log2 fold change for differentially expressed genes in each cluster that fall into the Meiosis I GO category (GO: 0007127). *p<0.05, **p<0.01, ***p<0.001, two-sided one-sample t-test compared to an expected value of 0.

*Smarca5* mRNA was predominantly expressed in spermatogonial populations, beginning early in spermatogenic development and continuing to the stage immediately preceding the major transcriptional and morphological defects observed in *Smarca5* cKO testes (**Fig 4D**). Low-moderate expression of *Smarca5* transcript continues through the end of meiosis, and was significantly downregulated in most of the corresponding cell clusters in the cKO condition (**Fig 4E**). Examination of differential gene expression between control and cKO in each cluster revealed that transcriptional changes begin at the undifferentiated spermatogonia stage (**Fig 4F, Supplemental Tables S1 and S2**). Differentially expressed genes (DEGs) were biased toward downregulation prior to meiosis and upregulation during meiotic prophase, while after meiosis many more transcriptional changes occur in both directions, which may be due to indirect regulation or secondary effects of meiotic failure. Both up- and down-regulated DEGs were enriched for functional categories related to spermatogenesis and cell cycle regulation (**Fig 4G**). Upregulated DEGs additionally were enriched for functions related to meiotic prophase, such as homologous recombination, chromosome organization, and piRNA processes. In contrast, downregulated genes were more enriched for functions related to spermatid differentiation, such as flagellated sperm motility and spermatid development. Genes associated with meiosis I (GO:0007127) were mildy but significantly downregulated prior to meiosis and then upregulated in late meiotic and postmeiotic stages, suggesting defective activation and developmental timing of expression of genes required for meiotic progression (**Fig 4H**).

We next assessed changes in chromatin accessibility induced by loss of SMARCA5. We isolated control and *Smarca5* cKO differentiating (cKIT+) spermatogonia by flow cytometry and collected ATAC-seq data from two biological replicates of each genotype (**Supplemental Fig S4A, Supplemental Tables S3 and S4**). cKIT+ spermatogonia represent a developmental time point coinciding with the peak of SMARCA5 mRNA expression and immediately preceding the onset of major phenotypic defects observed in the *Smarca5* cKO. Therefore, we anticipated that chromatin changes in these cells were most likely to be direct effects of SMARCA5 loss rather than secondary to other cellular or tissue defects. We observed a global increase in accessibility at transcription start sites (TSS) in *Smarca5* cKO germ cells (**Fig 5A, 5B**). While there were some peaks gained in the *Smarca5* cKO condition, the majority of peaks found in the cKO were also found in control (**Fig 5C, Supplemental Tables S5 and S6**), suggesting that loss of SMARCA5 permits additional chromatin opening at sites that are already accessible in wild type cells. Consistent with this model, clustering based on ATAC-seq signal revealed that among TSS with strong (cluster 1) or moderate (cluster 2) gains in accessibility, most were already at least partially accessible in control. On the other hand, inaccessible TSS largely remained inaccessible in *Smarca5* cKO cells (**Fig 5D**). Interestingly, genes whose expression was upregulated in the differentiating spermatogonia/meiotic entry cluster (**Fig 4**) displayed increased accessibility centered on the TSS as well as at upstream nucleosomal regions (**Fig 5E**). Genes whose expression was downregulated in differentiating spermatogonia also gained accessibility, but this was limited to the TSS region (**Fig 5E**). Among the upregulated DEGs that gained TSS accessibility was *Sycp3*, a component of the synaptonemal complex whose overexpression could contribute to the synapsis defects observed in meiotic spreads (**Fig 5F**). Additional genes important for meiosis and transcriptionally misregulated at either the differentiating spermatogonia or early pachytene stages also showed gains in accessibility (**Fig 5F, Supplemental Fig S4B**). We conclude that loss of SMARCA5 substantially disrupts normal chromatin remodeling during the early stages of male meiotic prophase, leading to aberrant gene expression, failure to generate normal post-meiotic germ cells, and sterility.

**Figure 5.**
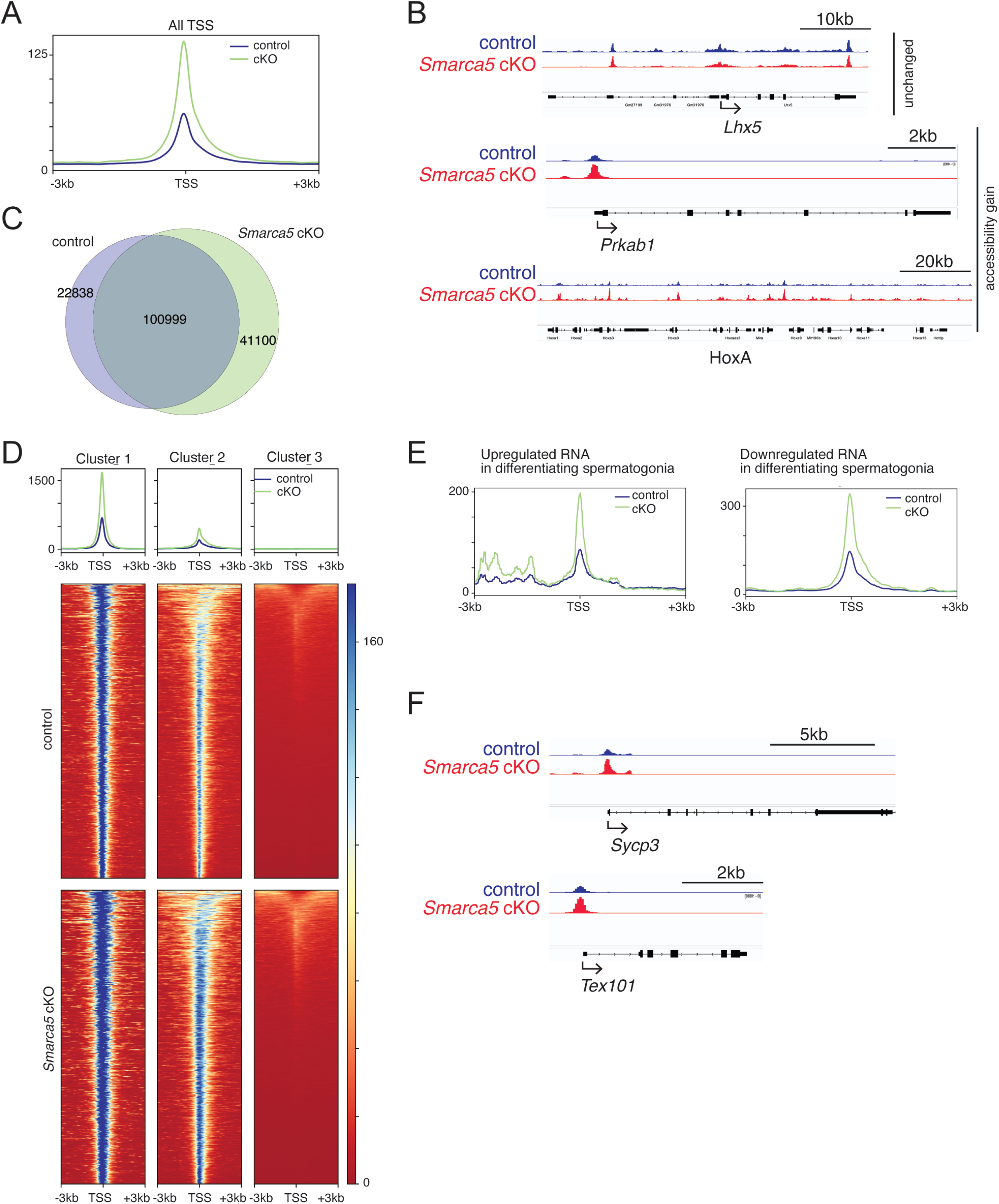
Loss of *Smarca5* leads to increase in chromatin accessibility. **A**, Metagene plot showing ATAC-seq signal at all transcription start sites (TSSs) in control and *Smarca5* cKO differentiating spermatogonia. **B**, Genome browser snapshots showing ATAC-seq signal in control and *Smarca5* cKO differentiating spermatogonia. *Lhx5* is an example of a gene with no change in accessibility; *Prkab1* and the HoxA cluster show gains in accessibility. **C**, Overlap between ATAC-seq peaks called in control and *Smarca5* cKO germ cells. **D**, Clustering of ATAC-seq signal at TSS showing strong gain in accessibility (cluster 1), moderate gain in accessibility (cluster 2) and no change (cluster 3). **E**, Metagene plots showing ATAC-seq signal at genes whose expression is up- or down-regulated in the differentiating spermatogonia (see **Fig 4**). **F**, Genome browser snapshots at *Sycp3* and *Tex101*, genes whose expression is upregulated in *Smarca5* cKO differentiating spermatogonia.

Regulation of meiosis is important for formation of functional gametes, and meiotic defects are a leading cause of infertility in both sexes. We found that the chromatin remodeler SMARCA5 is essential for meiotic progression in mammalian male germ cells. SMARCA5 regulates meiosis in spermatogenesis by maintaining a restricted chromatin architecture to ensure appropriate timing and levels of gene expression, and its loss leads to aberrantly high chromatin accessibility across the genome and widespread transcriptional defects. This effect contrasts with its role in female germ cells, where SMARCA5 acts to open chromatin and promote meiotic gene expression. This difference in function between the sexes may relate to the differences in male and female nuclear states during the relevant stages of meiotic prophase, since growing and fully-grown germinal vesicle (GV) oocytes are highly differentiated, while spermatogonia may have a more flexible, precursor-like chromatin architecture even as they approach meiotic entry. Interestingly, we also observed elevated expression of the LINE1 retroelement in *Smarca5* cKO spermatocytes, suggesting that aberrant chromatin opening in the absence of SMARCA5 may permit unlicensed derepression of retrotransposons. SMARCA5 thus may be important for shaping the meiotic transcriptome in males by guiding transcriptional activity towards required genes and restricting promiscuous expression from other loci.

## Materials and Methods

### Mice

All mice were maintained and euthanized under standard conditions according to the principles and procedures described in the National Institutes of Health Guide for the Care and Use of Laboratory Animals. These studies were approved by the Yale University Institutional Animal Care and Use Committee under protocol 2023-20169. *Smarca5* cKO mice were generated by crossing *Smarca5* ^*fl/fl*^ mice (Alvarez-Saavedra et al. 2014) with *Ddx4-Cre* (Gallardo et al. 2007; Hu et al. 2013) or *Spo11-Cre* (Lyndaker et al. 2013). For fertility testing, *Smarca5* cKO or littermate control (*Cre*-negative, *Smarca5*^*fl/+*^) males were co-housed with wild type females for six months starting at three months of age and litter sizes were recorded at weaning.

### Histology

Testes and epididymides were dissected at 3-4 months of age. For *Smarca5;Spo11-Cre*, a heterozygous littermate control was used instead of the standard control. After isolation, tissues were briefly washed with phosphate-buffered saline (PBS) and fixed in Hartman’s fixative (Millipore-Sigma; H0290-500ML) for 48 h at room temperature (RT). Samples were then washed in 70% ethanol, dehydrated and embedded in paraffin wax. 4 µm thick sections were prepared on glass slides, cleared in xylene and dehydrated in a graded series of ethanol. The sections were then stained with hematoxylin and eosin (H&E). H&E images were acquired with a bright-field microscope (Zeiss Axio Lab A1 Bright Field Microscope, 20×0.5 NA or 40×0.75 NA objectives).

### Immunofluorescence staining

Paraffin wax-embedded testis sections were deparaffinized in xylene, dehydrated in a graded series of ethanol, boiled in 10 mM sodium citrate buffer (pH 6) in a microwave oven for 20 min to retrieve the antigen, washed in PBS, permeabilized in 0.5% Triton X-100 for 10 min, blocked in blocking buffer (5% BSA+0.1% Triton X-100) for 1 h at RT and then incubated overnight at 4°C with primary antibodies diluted in blocking buffer. Slides were washed with PBS and stained with fluorophore-conjugated secondary antibodies, diluted at 1:500 in blocking buffer and incubated at RT for 1 h in the dark. Slides were counterstained with Hoechst 33342 (Thermo Fisher Scientific) for 5 minutes at RT in the dark and finally mounted in Antifade mounting medium (Vector Labs). Images were acquired with an LSM 980 airyscan confocal microscope (Zeiss) or a Stellaris DIVE (Leica) microscope equipped with 405, 488 or 555/561 nm lasers, and fitted with a 63×1.4 NA objective. Images were processed with Zen acquisition software and ImageJ. Catalog numbers and dilutions of primary and secondary antibodies used in this study are listed in the table below.

### Preparation of testicular cell suspensions for flow cytometry

Testicular cells from adult (3 month old) *Smarca5* cKO male mice and littermate (*Cre*-negative, *Smarca5*^*fl/+*^) controls were prepared for flow cytometry analysis as described previously (Malla et al. 2023). Testes were dissected, the tunica albuginea was removed, and the seminiferous tubules were minced in Ca^2+^- and Mg^2+^-free PBS (Gibco). Cells were dispersed by gentle aspiration, filtered using a 40 µm nylon filter, and washed in PBS by centrifuging at 800×*g* for 5 min. The cells were re-suspended in PBS, fixed in 70% chilled ethanol, and stored at 4°C for 24 h or at −20°C for up to 1 week until further analysis. Immediately before analysis, 1×10^6^-2×10^6^ ethanol-fixed testicular cells were washed three times with PBS and treated with 0.25% pepsin solution for 10 min at 37°C. Finally, cells were stained with propidium iodide (PI) staining solution (25 µg/ml PI, 40 mg/ml RNase A and 0.03% Nonidet P-40 in PBS) at RT for 20 min. The PI-stained cells were analyzed on Bio-Rad S3e cell sorter (Bio-Rad; excitation 488 nm; emission 585/40 nm) as described previously (Krishnamurthy et al. 2000). Analysis was performed using FlowJo software, with gates set based on SSC and PI-A and applied identically between control and *Smarca5* cKO samples.

### Preparation and staining of meiotic spermatocyte spreads

Meiotic chromosome spreads were prepared as described previously (Peters et al. 1997; Malla et al. 2023). Testes from young adult (3 month old) *Smarca5* cKO and littermate control male mice were dissected in cold PBS, and the tunica albuginea and extracellular material were removed. After two quick washes in cold PBS, the seminiferous tubules were incubated in hypotonic extraction buffer [30 mM Tris (pH 8.2), 50 mM sucrose, 17 mM trisodium citrate dihydrate, 5 mM EDTA, 0.5 mM dithiothreitol and 0.5 mM phenylmethylsulfonyl fluoride] containing 1X protease inhibitor cocktail (Roche, 11836153001) for 1 h at RT. The tubules were then removed from the hypotonic buffer, transferred to a glass slide, and minced to release the cells. 10 μl of cell suspension was diluted with 40 μl 100 mM sucrose and spread onto a glass slide pre-dipped in 1% paraformaldehyde (PFA) containing 0.15% Triton X-100. Slides were dried for 6 h in a humidified chamber before proceeding with immunofluorescence staining. After two washes in PBS, the dried slides were treated with 2% PFA and 0.15% Triton X-100 for 10 min at RT and then blocked in blocking buffer (5% BSA+0.1% Triton X-100) for 1 h at RT. Slides were then incubated with primary antibody diluted in blocking buffer overnight at 4°C. After three washes in PBS, slides were stained with fluorophore-conjugated secondary antibodies. All secondary antibodies were used at 1:500 dilution in blocking buffer and incubated at RT for 1 h. Slides were counterstained with Hoechst 33342 (Thermo Fisher Scientific) for 5 minutes at RT in the dark and finally mounted in antifade mounting medium (Vector Labs). Images were captured with LSM 980 airyscan confocal microscope (Zeiss) or Stellaris DIVE (Leica) microscope equipped with 405, 488 or 555/561 nm lasers, and fitted with a 63×1.4 NA objective. Images were processed with Zen acquisition software and ImageJ.

### TUNEL assay

TUNEL assay was performed on paraffin embedded sections using the Abcam TUNEL Assay HRP-DAB kit (ab206386) as per the manufacturer’s protocol.

### Sorting of cKIT-positive spermatogonia and ATAC-seq

A polypropylene collection tube was coated with 3 ml Collection Buffer (20% FBS in 1X PBS) for 2 hours at RT with end-over-end mixing. Seminiferous tubules were mechanically isolated from the tunica and transferred to a 15ml conical tube containing 5ml Digestion Solution I [0.75 mg/ml collagenase type IV (Gibco, 17104-019) in Dulbecco’s modified Eagle’s media (DMEM, Gibco, 11965092) supplemented with 1:1000 with 1 mg/ml DNase I (StemCell Technologies, 07900)] warmed to 37 °C, followed by incubation at 37 °C for 10 mins with end-over-end rotation. 5ml of cold DMEM was added and tubules allowed to settle on ice. The supernatant was discarded and the tubules were washed again with 5ml cold DMEM to deplete Leydig cells. 5ml of DMEM was added followed by centrifugation at 500xg for 5 mins at 4°C. The supernatant was removed and 5ml of Digestion Solution II [Accutase (Gibco, A1110501) supplemented with 1 mg/ml DNase I at 1:1000 at room temperature)] was added to the tubules followed by incubation with end-over-end rotation for 10 mins at RT with gentle pipetting after 5 mins to disperse the tubules. 5ml of cold Complete Media (10% FBS in DMEM) was added and the sample was centrifuged at 500 xg for 5 mins at 4 °C. The cell pellet was then resuspended in 5 ml of cold Complete Medium, filtered through a 100 μm strainer, and the cell suspension was transferred to a fresh 15 ml conical tube and centrifuged at 500 xg for 5 mins at 4 °C. The pellet was resuspended in 1 ml of cold Complete Medium with 1 μl (0.2 µg) of PE-conjugated anti-KIT antibody and incubated for 20 mins on ice protected from light with occasional mixing. An aliquot of unstained cells (∼ 0.5 ml) was reserved for gating. 4 ml cold Complete Medium was added to the tube followed by centrifugation at 500 xg for 5 mins at 4°C. The cell pellet was washed twice with 5 ml of cold Complete Medium and resuspended in 3 ml of cold FACS buffer (5% FBS in 1X PBS). Cells were filtered through a 70 µm strainer and transferred to a 5 ml polypropylene round-bottom tube on ice. For sorting, the coated collection tube was filled with 1 ml Collection Buffer. Samples were gated to remove debris and select singlets, and cKIT+ cells were gated relative to the unstained aliquot in the FL2 channel.

### Sequencing and data analysis for ATAC-seq

Approximately 50,000 sorted cKIT+ cells were processed for each ATAC-seq library using the Active Motif kit (53150) as per the manufacturer’s protocol. ATAC-seq libraries were sequenced on an Illumina NovaSeq instrument at a read depth of 30 million paired-end reads per library. Low quality reads were filtered and the adaptors were trimmed using Cutadapt (Martin 2011). ATAC-seq reads were aligned to the mm10 reference genome using bowtie2 (Langmead and Salzberg 2012) in very sensitive mode. Reads aligning to mitochondrial DNA were discarded and PCR duplicates were removed using Picard tools (https://broadinstitute.github.io/picard). Bigwig tracks were generated using bamcoverage from the deeptools package and peaks were called using MACS2 (Zhang et al. 2008) with the –broadpeak option. After confirming a high correlation between replicates, replicate peaks were combined using the merge function in BEDTools (Quinlan and Hall 2010). Multibamsummary and computematrix from deeptools (Ramirez et al. 2014) was used for further analysis. Peaks were assigned to the single nearest gene within 1000 base pairs using the Genomic Regions Enrichment of Annotations Tool (GREAT).

### Single cell RNA sequencing

15,000 dissociated *Smarca5* cKO testis cells (“cKO”) were processed using a Chromium Next GEM Single Cell 3’ v3.1 kit (10X Genomics) and sequenced using an Illumina NovaSeq machine with 150 base pair paired-end reads at a depth of 250 million reads. A previously published wild type dataset (“control”) generated by our lab under identical conditions (GSE216343: GSM6670717, GSM6670718, and GSM8289596) was used for comparison.

Sequences were transformed into raw count matrices based on an mm10 reference using CellRanger (10X Genomics) and loaded into an R environment (R 2023) with Seurat 4.1.0 (Hao et al. 2021). The SoupX pipeline (Young and Behjati 2020) was used to remove ambient RNA contaminants from both datasets. These datasets were transformed into Seurat objects, merged, and filtered to remove doublets or multiplets (nCount_RNA < 25,000), dead or dying cells (nFeature_RNA < 7,000), or cells with high mitochondrial content (> 10%). After normalizing and scaling, ElbowPlot() was used to determine optimal dimensionality (19 dimensions). PCA was used as a method for linear dimensional reduction. Clusters were identified at a resolution of 0.5, RunUMAP() was used to generate UMAP objects, and clusters were further defined using key spermatogenic genes (Malla, Rainsford 2023; Walters, Rainsford 2024). Three clusters were removed from further analysis due to dead or dying cells, and ambiguous match to known testicular cell populations. The resulting dataset was normalized, scaled, and dimensionally reduced. A new UMAP was created at a resolution of 0.5, and integrated using Harmony (Korsunsky et al. 2019. The final UMAP was generated using the integrated dataset with 17 dimensions at a 0.5 resolution. Clusters were defined again using key spermatogenic or testicular somatic cell markers, and differentially expressed genes were identified by the FindMarkers() function. Graphs were created in R using the ggplot2 package {Wickham, 2016 #35). Gene ontology analysis was performed in R using the GOStats package (Falcon and Gentleman 2007).

## Data availability

Single cell RNA-seq and ATAC-seq data generated for this study is available at the NCBI GEO repository under accession number GSE279216. Control scRNA-seq data has been previously published under accession number GSE216343 (GSM6670717, GSM6670718, and GSM8289596).

## Antibodies used in this study

| Antibody | Company/Cat. # | Application | Dilution |
| --- | --- | --- | --- |
| SYCP3 | Abcam, ab97672 | IF | 1:300 |
| MVH | Cell Signaling Technology, 8761S | IF | 1:400 |
| SMARCA5 | Abcam, ab3749 | IF | 1:300 |
| SMARCA5 | Thermo Scientific PA5-52601 | WB | 1:1000 |
| Gamma H2AX | Abcam, ab2893 | IF | 1:400 |
| Gamma H2AX | Millipore, 05-636 | IF | 1:400 |
| SYCP1 | Abcam, ab15090 | IF | 1:400 |
| GAPDH | Santa Cruz, sc32233 | WB | 1:200 |
| LINE1 | Abcam, ab216324 | IF | 1:400 |
| Rabbit IgG (Alexa Fluor 488) | Invitrogen, A11034 | IF | 1:500 |
| Mouse IgG (Alexa Fluor 568) | Invitrogen, A11031 | IF | 1:500 |

## Competing interest statement

The authors declare no competing interests.

## Acknowledgments

We thank David Picketts and Arthur Skoultchi for the gift of *Smarca5*^*fl/fl*^ mice, and Paula Cohen for the gift of *Spo11-Cre* mice. We thank Benjamin Walters and Haoming Yu for help with sorting. We are grateful to the Yale Center for Genome Analysis, the Yale Flow Cytometry Facility, and the Yale Center for Cellular and Molecular Imaging for resources and assistance. SK was funded by the Surdna Foundation and Yale Venture Fund Fellowship. This work was supported by the National Institute of Child Health and Human Development (R01HD098128 and R21HD110843 to BJL). BJL is a Pew Scholar, supported by the Pew Charitable Trust.

## Author contributions

Conceptualization: SK, BJL; Investigation: SK, ABM, SRR; Validation: SK, ABM, SRR; Formal analysis: SK, ABM, SRR, BJL; Visualization: SK, BJL; Writing – original draft: SK; Writing – review & editing: BJL; Resources: BJL; Supervision: BJL; Funding acquisition: BJL

## Supplemental Material

**Supplemental Figure S1. Validation of *Smarca5* cKO mice. A**, Design of the *Smarca5* conditional allele and allele resulting from Cre excision (Alvarez-Saavedra et al. 2014). **B**, Genetic cross to generate *Smarca5* cKO males. **C**, Oocytes generated from *Smarca5* cKO females following super-ovulation. Left, example clutches from a single female. Right, quantitation from n=3 animals. *p<0.05, **p<0.01, ***p<0.001, unpaired t-test. **D**, Representative low-magnification images of γH2A.X staining in control and cKO testes. Scale bar, 50µm.

**Supplemental Figure S2. Defective spermatogenesis following conditional knockout of *Smarca5* in early meiotic prophase using *Spo11-Cre*. A**, Western blot for SMARCA5 in wild type and *Smarca5* conditional knockout (cKO) adult whole testes where cKO is driven by either *Ddx4-Cre* (middle lanes) or *Spo11-Cre* (right lanes). GAPDH is a loading control. Arrow indicates expected size for SMARCA5. **B**, Fertility of control and *Smarca5* cKO males (n=3-4). Each dot represents mean litter size for one male. Error bars represent standard deviation. No pups were born from *Smarca5;Spo11-Cre* cKO males. *p<0.05, **p<0.01, ***p<0.01, unpaired Welch’s t-test. **C**, Left, gross image of whole testis from control, *Smarca5;Spo11-Cre* heterozygous and *Smarca5;Spo11-Cre* cKO adult male mice. Each tick on the ruler represents 1mm. Right, testicular weight relative to body weight plotted for n=2-3 control, heterozygous and cKO testes. The same mice were plotted in the control condition for both *Ddx4-Cre* and *Spo11-Cre* comparisons (see **Fig 1D**). *p<0.05 **p<0.01 ***p<0.001, unpaired t-test. **D-E**, Hematoxylin and eosin (H&E) staining of paraffin-embedded sections from *Smarca5;Spo11-Cre* heterozygous and cKO adult testis (**D**) and epididymis (**E**). Scale bar, 50 µm. **F**, Representative images (left) and quantitation (right) of γH2A.X staining in *Spo11-Cre* heterozygous and cKO testes. Scale bar, 20µm. *p<0.05, unpaired t-test.

**Supplemental Figure S3. Quality control and validation for scRNA-seq data. A**, Feature counts, transcript counts, and mitochondrial reads for original datasets before filtering and Harmony. **B**, First two principal components for WT and cKO after Harmony. **C**, Elbow plot after Harmony. **D**, UMAP plot after Harmony showing WT and cKO distributions. **E**, Representative expression markers selected from the sets used to assign identities to each cluster projected onto control UMAP plots.

**Supplemental Figure S4. ATAC-seq in cKIT+ differentiating spermatogonia. A**, Scatter plots showing correlation in ATAC-seq signal between biological replicates. **B**, Genome browser snapshot showing ATAC-seq signal at *Sycp1*, a DEG downregulated in early pachytene cells that gains accessibility in differentiating spermatogonia.

**Supplemental Table S1. Upregulated differentially expressed genes**.

**Supplemental Table S2. Downregulated differentially expressed genes**.

**Supplemental Table S3. Control ATAC-seq peaks, merged replicates**.

**Supplemental Table S4. *Smarca5* cKO ATAC-seq peaks, merged replicates**.

**Supplemental Table S5. Control-only ATAC-seq peaks**.

**Supplemental Table S6. *Smarca5* cKO-only ATAC-seq peaks**.

